# Highly defended nudibranchs ‘escape’ to visually distinct background habitats

**DOI:** 10.1101/2022.12.13.520332

**Authors:** Cedric P. van den Berg, Matteo Santon, John A. Endler, Karen L. Cheney

**Author notes:** Corresponding author: Cedric P. van den Berg.

## Abstract

The ‘escape and radiate’ hypothesis predicts that once species have evolved aposematism, defended species can utilise more visually diverse visual backgrounds as they ‘escape’ the need to be well camouflaged. This enables species to explore new ecological niches, resulting in increased diversification rates. To test this hypothesis’ ‘escape’ component, we examined whether the background habitats of 12 nudibranch mollusc species differed among species depending on the presence and strength of chemical defences. We obtained a rich array of colour pattern statistics using Quantitative Colour Pattern Analysis (QCPA) to analyse backgrounds viewed through the eyes of a potential predator (triggerfish, *Rhinecanthus aculeatus*). Colour pattern analysis was done at viewing distances simulating an escalating predation sequence. We identified four latent factors comprising 17 non-correlated colour pattern parameters, which captured the among-species variability associated with differences in chemical defences. We found that chemically defended species, indeed, were found on visually distinct backgrounds with increased colour and luminance contrast, independent of viewing distance. However, we found no evidence for increased among-species background diversity coinciding with the presence and strength of chemical defences. Our results agree with the ‘escape and radiate’ hypothesis, suggesting that potent chemical defences in Dorid nudibranchs coincide with spatiochromatic differences of visual background habitats perceived by potential predators.

**Lay Summary:** According to the ‘escape and radiate’ hypothesis, the ability to store potent chemical defences as protection from predators enables animals to be minimally dependent on matching visual backgrounds for camouflage. We found evidence supporting this hypothesis in 12 species of nudibranch molluscs. Aposematic species were found on visual backgrounds that differed consistently in appearance from the visual background of undefended cryptic species. However, we found no difference in the variability of visual backgrounds between species with or without chemical defences. This suggests that warning colouration in eastern Australian Dorid nudibranchs coincides with broadly generalisable spatiochromatic properties of visual backgrounds.

## Introduction

Animals and plants use aposematic colour patterns to advertise defences to potential predators (Poulton, 1890). One of the hypothesised benefits of an organism evolving aposematism is the potential to ‘escape’ the costs of crypsis, including limited movement and thus access to resources due to the need to match a specific background habitat or preventing detection through motion contrast for primary defence (Endler, 1984; Merilaita & Tullberg, 2005; Regan & Beverley, 1984). Therefore, Merilaita and Tullberg (2005) suggested that alternative visual defence strategies, such as aposematism, are more likely to evolve in highly variable visual environments where crypsis is difficult to achieve. The evolution of aposematic signalling provides increased access to resources across ecological niches as individuals are not strongly restricted to particular habitats to avoid predator detection. This reduced dependency on crypsis may drive increased speciation and diversification rates (Arbuckle & Speed, 2015; Ehrlich & Raven, 1964). This process was coined ‘escape and radiate’ by Thompson (1989) and has since been adopted as a core concept underpinning the role of chemical defences in adaptive radiations (e.g. Arbuckle & Harris, 2021).

The evolution of potent chemical defences often coincides with aposematism (see (Ruxton et al., 2018; Summers et al., 2015; White & Umbers, 2021) for reviews), and theoretical modelling supports the conclusion that aposematic species are less constrained by the visual properties of their habitat (habitat generalists) than undefended background-matching species (habitat specialists) (Merilaita & Tullberg, 2005; Speed, Brockhurst, et al., 2010). The appearance of an animal against its visual background fundamentally impacts its detectability by potential predators, a crucial factor shaping the design and function of defensive colouration (see van den Berg, Endler, et al., 2023) for discussion). Therefore, assuming relaxed selection for background matching, secondary defences in a diverse prey community inhabiting visually complex habitats might facilitate chemically defended species to inhabit distinct and possibly more diverse visual backgrounds than undefended species. However, whether the presence of chemical defences qualitatively or quantitatively correlates with general among-species differences in background habitats remains unknown.

Here, we tested the hypothesis that more defended species inhabit distinct and more variable habitats using 12 species of co-occurring Eastern Australian nudibranch species (Fig. S1). Nudibranchs are an intriguing system for the study of defensive animal colouration due to potent chemical defences in many species (Avila, 1995; Winters et al., 2022; Winters, White, et al., 2018) and colour patterns ranging and colour patterns that range from bold aposematic displays to near-perfect camouflage (Debelius & Kuiter, 2007). Indeed, the evolution of specialised glands (mantle dermal formations) for the storage and secretion of defensive chemicals (Carbone et al., 2013; Wägele et al., 2006), in combination with the evolution of aposematic signalling, may have led to adaptive radiation in nudibranchs (Gosliner, 2001).

Until recently, studies quantifying visual backgrounds in the context of defensive animal colouration have relied on the independent consideration of spatial and spectral (colour and luminance) properties (e.g. Cortesi & Cheney, 2010; Nokelainen et al., 2021). In this study, we considered these pattern components in combination (van den Berg, Troscianko, et al., 2020) using calibrated digital photography, the multispectral image calibration and analysis toolbox (MICA) (Troscianko & Stevens, 2015) and its’ integrated Quantitative Colour Pattern Analysis (QCPA) framework (van den Berg, Troscianko, et al., 2020). This methodology enabled us to assess the spatiochromatic information of natural backgrounds upon which nudibranch individuals (12 species, n = 184) were found. QCPA provides a descriptive array of image statistics capturing the spatiochromatic properties of each background according to the physiological limitations of an ecologically relevant observer. Here, we used the visual system of a triggerfish (*Rhinecanthus aculeatus*), a common, omnivorous reef fish. We analysed different viewing distances to model an escalating predation sequence to account for changes in background appearance to a predator up close (2cm) and at a distance (30cm).

We then compared these colour pattern parameters to measures of the strength of chemical defence for each species, which ranged from highly unpalatable to palatable, using previously published data from antifeedant assays with rockpool shrimp (*Palaemon serenus*) and brine shrimp (Winters et al., 2022). We hypothesised that chemically defended (unpalatable) species would be found on visually distinct, more variable visual backgrounds than undefended (palatable) species. Furthermore, we predicted a strong correlation between the presence and strength of chemical defences and differences in spatiochromatic properties of background habitats among species with differing strengths of chemical defences at distances where prey detection and identification are most likely (Endler, 1986, 1991; Ruxton et al., 2018).

## Materials and Methods

### Study species

We took calibrated digital photographs of 184 Dorid nudibranch individuals belonging to 12 species from four locations on the east coast of Australia: Sunshine Coast (SE Queensland, QLD), Gold Coast (SE QLD), Cook Island (New South Wales, NSW) and Nelson Bay (NSW) between March 2016 and February 2021 (Table S1, Fig. S1). Our study considers many of the more commonly found Dorid nudibranchs in the study sites (e.g. (Larkin et al., 2018; Schubert & Smith, 2020; Smith & Davis, 2019)). Species were identified visually using various nudibranch ID guides (Coleman et al., 2015; Debelius & Kuiter, 2007; Gosliner et al., 2018): *Aphelodoris varia* (n = 22), *Chromodoris elisabethina* (n = 21), *Chromodoris kuiteri* (n = 17), *Chromodoris lochi* (n = 3), *Dendrodoris krusensterni* (n = 7), *Discodoris sp.* (n = 13), *Doriprismatica atromarginata* (n = 27), *Glossodoris vespa* (n = 15); *Hypselodoris bennetti* (n = 10); *Phyllidia ocellata* (n = 23); *Phyllidia varicosa* (n = 8); *Phyllidiella pustulosa* (n = 18); numbers indicate sample sizes used in the analysis. *Discodoris sp*. in our study may have constituted a mixture of closely related species, including *Sebadoris fragilis*, *Jorunna pantheris, Tayuva lilacina* and undescribed species; however, these species are visually indiscriminable. Therefore, they are named *Discodoris sp.* as they were found in the same locations, have no known chemical defences, are predominantly nocturnal and are closely related (Larkin et al., 2018). Only one out of the 12 species (*Doriprismatica atromarginata*) was found in comparable numbers across all sites in NSW and SE QLD, with most species found either in NSW or SE QLD (Table S1). Nudibranchs were photographed underwater against their natural habitat using a calibrated Olympus EPL-5 with a 60mm macro lens in an Olympus PT-EP10 underwater housing using white LED illumination from a combination of VK6r and PV62 Scubalamp video lights (van den Berg, Troscianko, et al., 2020). All pictures were taken at roughly a 90-degree angle relative to each animal and its background (Fig. 1).

**Figure 1.**
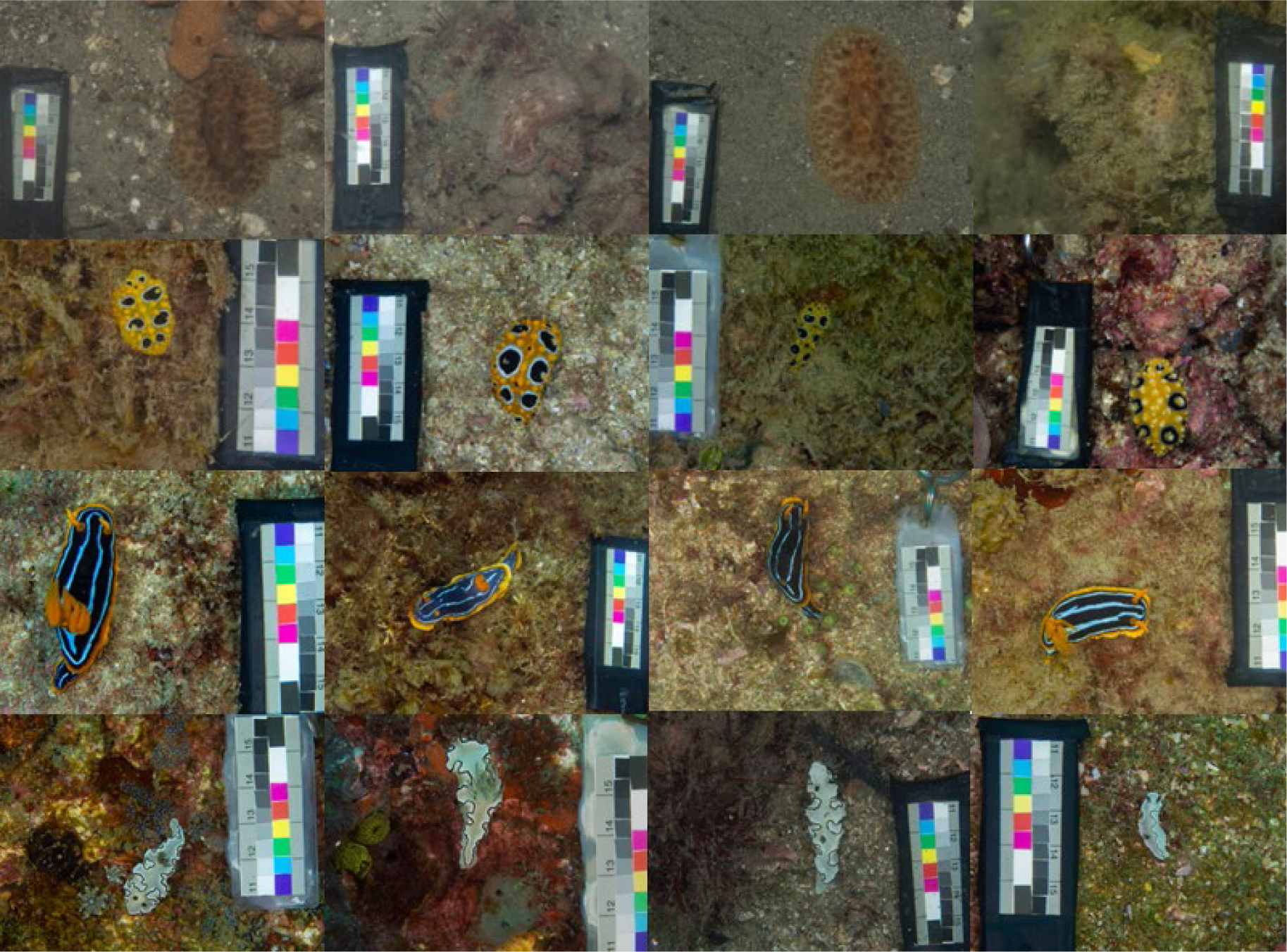
Representative nudibranch and background images for 4 out of the 12 species in this study. From top to bottom: *Discodoris sp., Phyllidia ocellata, Chromodoris kuiteri, Doriprismatica atromarginata*.

### Image analysis

Using ImageJ (Schneider et al., 2012) and the MICA toolbox (Troscianko & Stevens, 2015), the images were manually segmented into regions of interest (ROI), selecting the animal from its background and defining a size standard using a custom-made colour and grey standard placed in each image (Fig. 1). All images were aligned, so the nudibranch was aligned so the majority of each animal’s body was oriented vertically (head up) before analysis using QCPA (van den Berg, Troscianko, et al., 2020). To analyse the images, we used the visual system parameters of a trichromatic triggerfish, *Rhinecanthus aculeatus* (with spectral sensitivities of 413 nm, 480 nm and 528 nm λ_max_; Cheney et al. 2022). This species (adult total length, TL ∼ 15 cm) is a typical shallow reef inhabitant found throughout the Indo-Pacific and feeds on invertebrates, algae, and detritus (Randall et al., 1997) and is deemed a potential predator of nudibranchs. We analysed for viewing distances of 2 cm and 30 cm, assuming a triggerfish spatial acuity of 3 cycles per degree (Champ et al., 2014). The viewing distances reflect a range covering the maximal distance at which the largest specimen and coarse background detail would be detectable (30cm) to the closest possible distance, reflecting ultimate contact between predator and prey or visual background 2cm). Following acuity modelling, the images were processed with a Receptor Noise Limited (RNL) ranked filter (falloff: 3, radius: 5, repetition: 5) and clustered using RNL clustering with a colour discrimination threshold of 2 ΔS (Cheney et al., 2019) and a luminance contrast threshold of 4 ΔS (van den Berg, Hollenkamp, et al., 2020). We used Weber fractions based on a relative photoreceptor abundance of 1:2:2:2 (sw:mw:lw:dbl) and photoreceptor noise of 0.05, resulting in 0.07:0.05:0.05:0.05.

QCPA analysis was achieved using a custom batch script (van den Berg, Condon, et al., 2023), resulting in a highly descriptive array of 157 colour pattern statistics for each animal’s visual background. These parameters were spread across the following subtypes of colour pattern analysis: 1) Colour Adjacency analysis (CAA). CAA uses a transition matrix tallying all the synonymous and non-synonymous transitions along horizontal and vertical sampling transects across an image segmented into colour pattern elements. This transition matrix is then used to describe the geometry of a colour pattern (Endler, 2012; van den Berg, Troscianko, et al., 2020). 2) Visual contrast analysis (VCA). VCA uses the relative abundance and spectral properties of colour pattern elements to describe visual contrast (Endler & Mielke, 2005; van den Berg, Troscianko, et al., 2020). 3) Boundary strength analysis (BSA). BSA uses the relative abundance of boundaries between colour pattern elements to describe visual contrast (Endler et al., 2018; van den Berg, Troscianko, et al., 2020). 4) Local edge intensity analysis (LEIA). LEIA quantifies the strength and abundance of edge contrast in an unsegmented image. Statistics ending with ‘hrz’ or ‘vrt’ are the horizontal (across body axis) and vertical (along body axis) versions (analysing the respective transition matrix only) of their respective statistic (analysing the full transition matrix). For example, a background with algae or seagrass will likely have an elongated aspect ratio, whereas a uniform sandy background will not. A detailed description of each pattern statistic can be found in van den Berg, Troscianko, et al. (2020).

#### Level of chemical defence

As a measure of chemical defence for each species, we used previously published data from antifeedant assays with rockpool shrimp (*Palaemon serenus*) and toxicity assays with brine shrimp (Winters et al., 2022). Assay data for *Glossodoris vespa* is presented in Winters *et al*. (2018). In summary, the data were obtained from extracting secondary metabolites from nudibranchs and added to food pellets made from squid mantle at increasing concentrations. Effective dose (ED_50_) values were calculated based on the concentration that elicited a rejection response in at least 50% of the shrimp. Lethal dose (LD_50_) values were calculated based on the concentration that killed at least 50% of brine shrimp. For this study, the resulting ED_50_ and LD_50_ values were normalised and calculated as 1-ED_50_ / 1-LD_50_ to range from 0 (most palatable/toxic) to 1 (least palatable/toxic). Where multiple estimates of ED_50_ existed for a species due to multiple extracts/assays being performed, the average value was used. Only assays using whole-body extracts were considered to allow for comparisons between species.

Due to the uneven spread of toxicity and palatability values among species, we classified each species according to the shape of a sigmoidal response curve where species with values of 0 were considered palatable, values up to 0.25 were considered weakly unpalatable, values between 0.25 and 0.74 medium unpalatable and values of 0.74 and higher as highly unpalatable. We chose 0.74 rather than 0.75 as the boundary between medium and high unpalatability, as this was the median unpalatability of species with chemical defences (Table S1). To ensure at least three species in each group of chemical defences and to allow for subsequent investigations of differences in backgrounds between species with different levels of chemical fences, we separated the species into those who had no known toxicity and unpalatability: those with some level of toxicity and moderate unpalatability and those with some level of toxicity and high unpalatability (Fig S1).

## Statistical analysis

To analyse the large dataset derived from the QCPA analysis, we only kept images that did not produce any missing value for any background pattern metrics. VCA, CAA, and BSA metrics can produce NaN or infinite values if a colour pattern has less than two colour pattern elements following RNL clustering [60]. LEIA metrics do not suffer from this limitation. We then applied a Latent variable Exploratory Factor Analysis (EFA) with the R package *psych* using the factoring method of Ordinary Least Squares “ols” and the orthogonal rotation “varimax”. To prepare the dataset for the EFA, we first filtered the number of highly correlated QCPA metrics by keeping those that were less correlated than 0.6 Pearson correlation, which reduced their number from 157 to 17. We then ran the factor analysis based on four factors. The number of factors was selected by comparing the eigenvalues calculated from the original dataset to the median eigenvalues extracted from 10,000 randomly generated datasets with the same rows and columns as the original data. We selected factors with eigenvalues greater than the median of the eigenvalues from the simulated data. We also computed bootstrapped confidence intervals of the loadings by iterating the factor analysis 1000 times.

Due to data filtering for metrics less correlated than 0.6, the QCPA parameter listed for a given loading is likely synonymous with various other parameters in our 157-dimension colour pattern space. Therefore, the precise wording to describe each factor can vary substantially depending on which colour pattern metrics are put into focus (for a complete list of parameter correlations, see Table S2).

The scores of the factors extracted from the EFA were then used to implement four phylogenetic, distributional linear mixed models to compare the natural backgrounds of nudibranchs with different levels of chemical defences. Models were run in R v 4.1.2 (https://www.r-project.org/) using the brms package (Bürkner, 2018), which fits Bayesian models using Stan (https://mc-stan.org/). To account for the phylogenetic dependency of closely related species, all models included the phylogenetic tree of the 12 species of nudibranchs (Fig. S2), with the tree from (Cheney et al., 2014) pruned and missing species added according to their taxonomic classification in the World Register of Marine Species (Appeltans et al., 2012). The phylogenetic model was implemented following the guidelines of the online brms vignette (https://cran.rproject.org/web/packages/brms/vignettes/brms_phylogenetics.html) based on de Villemereuil & Nakagawa (de Villemereuil & Nakagawa, 2014).

The first model investigated differences in scores for latent factor 1 between nudibranchs with different levels of chemical defences (see chemical defences section) using a Student distribution. The model estimated the effect of the main categorial predictors level of chemical defence (low, moderate, high) and observer distance (2 cm and 30 cm) and their interaction on the response distribution’s mean and the residual standard deviation. To account for repeated measurements of each species, we also included species ID as a random intercept to the model. We further included random slopes over distance because their relationship with the value of the response factor 1 changes among species.

The second, third and fourth models were identical to the first but used factor 2, factor 3 and factor 4 as response variables. All models were fitted using weakly informative prior distributions (normal with mean = 0 and sd = 5 for intercept and coefficients, exponential (1) for standard deviations). Their performance was evaluated using posterior predictive model checking, which compares model predictions with observed data to assess overall model fit. We ran four Markov-Chain-Monte-Carlo (MCMC) chains for each model and obtained coefficient estimates from 16,000 post-warm-up samples. All model parameters reached reliable conversion indicators (Korner-Nievergelt et al., 2015): A Monte Carlo standard error smaller than 5% of the posterior standard deviation, an effective posterior sample size greater than 10% of the total sample size, and a *R̂* statistic value smaller than 1.01.

For graphical displays of the results, we present - for each combination of chemical defences and distance - the medians of latent factors values and their 95% Credible Intervals (CIs) of the posterior distributions of fitted values for the population average obtained from the joint posterior distributions of the model parameters (Korner-Nievergelt et al., 2015). The same posterior distribution of fitted values was used to compute pairwise differences and their 95% CIs between all possible combinations of the same two categorical predictors using the emmeans R package (Lenth, 2023). To compare the variance of response values between all combinations of predictor levels, we also compute the posterior distribution of all pairwise differences of the residual standard deviation on the original scale (back-transformed from the log scale). The effect size of pairwise differences increases with increasing deviation of such differences from zero, and the robustness of the result increases with decreasing degree of overlap of the 95% Credible Intervals (CIs) with zero.

## Results

Using the EFA, we identified four factors to describe how background pattern metrics differ between multiple viewing distances and levels of chemical defences. While not intended to identify a maximal amount of variability in colour pattern variation in our dataset, the four factors still explain 39% of the total variation (factor 1: 11%; factor 2: 11%, factor 3: 11%, factor 4: 6%) (Fig 2). Looking at the loadings of each factor, we can identify what latent variable they describe. While it would be possible to discuss each factor in almost infinite depth, we keep their description to loadings of +/-0.4 to capture their main properties.

**Figure 2.**
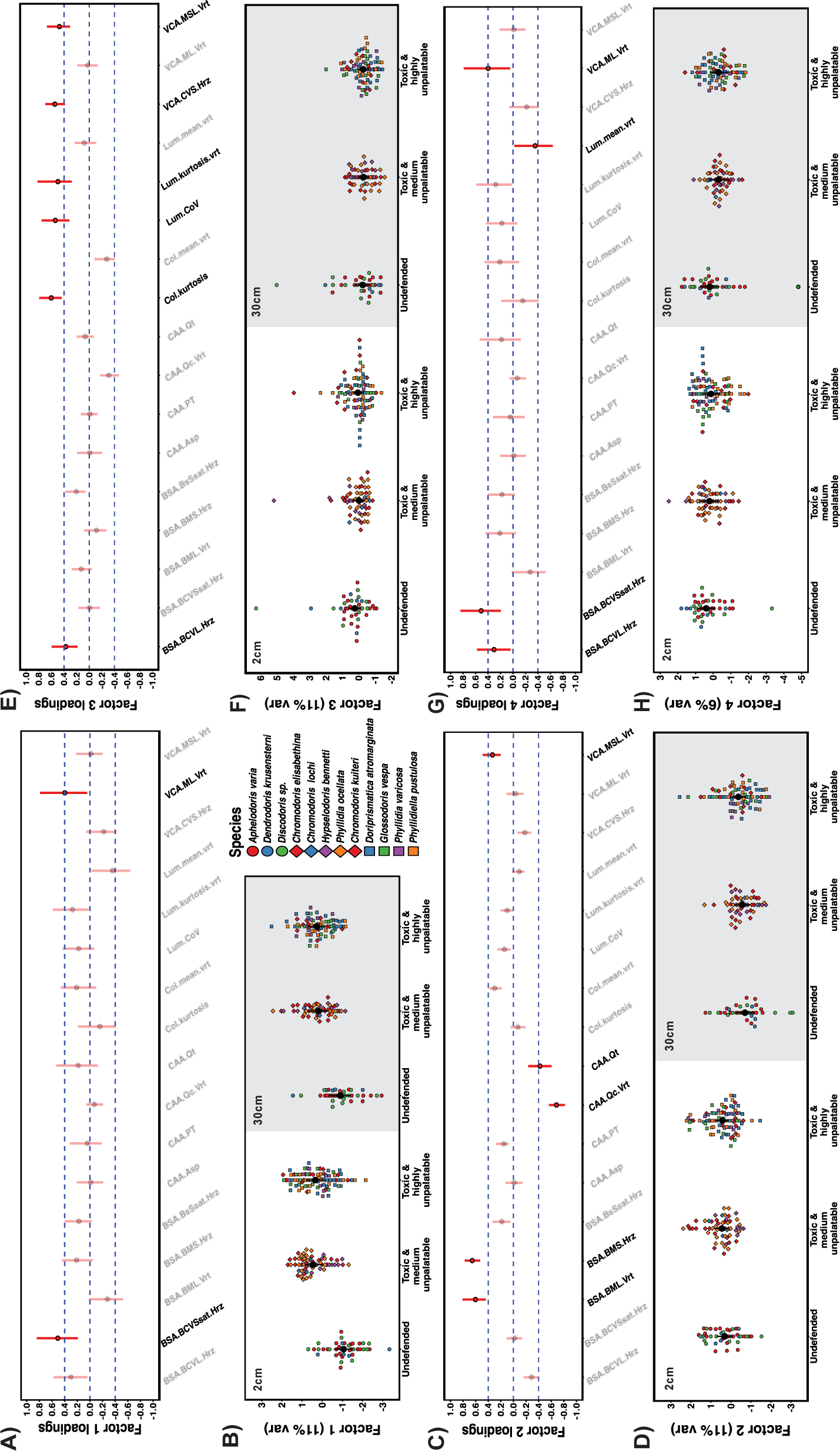
Detailed visual representation of the loadings of factor 1 (A), factor 2 (C), factor 3 (E), and factor 4 (G). Greyed-out factor loadings indicate colour pattern descriptors with minor contributions (<0.4) to each factor. Differences between group estimates of factor values are given for factor 1 (B), factor 2(D), factor 3 (F), and factor 4 (H). Estimates are given for 2cm viewing distance (left panel half, white) and 30cm (right panel half, grey).

### <u>Factor 1:</u> *Higher luminance and chromatic contrast coincides with smaller patch size*

Factor 1 (Fig. 2a) describes 11% of variation in visual backgrounds and captures a positive relationship between increases in the mean RNL luminance edge contrast (e.g., *Lum.mean*) and RNL chromaticity edge contrast (e.g. *Col.mean*) as measured by LEIA (unclustered image) and the average size of colour pattern elements in an RNL clustered image (*CAA.PT*). The increase in average luminance and chromatic edge contrast in visual backgrounds also coincides with an increase in variability of the luminance contrast relative to the mean (e.g. *BSA.BCVL*).

The visual backgrounds of chemically defended species have noticeably higher levels of factor 1 than those of undefended species [difference (+-95% CI)] (Fig 2a). This is true up close (2cm) as well as from further away (30cm) for toxic species with moderate levels of unpalatability (2cm: -1.53 (−2.28/ -0.78); 30cm: -1.05 (−1.83 / -0.23)) as well as toxic species with high levels of unpalatability (2cm: -1.44 (−2.18 / -0.73); 30cm -1.17 (−1.92 / -0.37)). However, toxic species with moderate levels of unpalatability have similar factor values to species with high levels of unpalatability up close (2cm: 0.08 (−0.38 / 0.54); 30cm: -0.13 (−0.63 / 0.36)). Therefore, chemically defended species were found on visual backgrounds with more chromatic and achromatic contrast (Table S3). However, the variability of backgrounds on which species were found did not differ between groups and remained similar at 2cm and 30cm (Table S4).

### <u>Factor 2:</u> *Higher colour and luminance contrast between patches coincide with lower background homogeneity*

Factor 2 (Fig. 2b) describes 11% of the variation in visual backgrounds and mostly captures the correlation between simultaneous increases in the strength of luminance contrast (e.g. *BSA.BML, VCA.MSL*) and saturation contrast (e.g. *BSA.BMS*) between colour pattern elements and decreases in background evenness (e.g. *CAA.Qt* & *CAA.Qc*).

We found no differences in factor values between species with different levels of chemical defences [difference (+-95% CI)] (Fig. 2b, Table S5) nor any differences between groups in the variability of factor values (Table S6) at 2cm or 30cm viewing distance. All groups showed higher factor values at 2cm compared to 30cm (Fig. 2b, undefended: 1.01 (0.57 / 1.48); toxic moderately unpalatable: 1.01 (0.67 / 1.31); toxic highly unpalatable: 0.80 (0.47 / 1.11)). The variability of backgrounds on which species were found did not differ between groups and remained similar at 2cm and 30cm (Table S6).

### <u>Factor 3:</u> *Increased luminance and colour contrast variability correlate with reduced average colour pattern regularity*

Factor 3 captures the relationship between increases in the variability of luminance contrast in visual backgrounds (e.g. *Lum.kurtosis, Lum.CoV*) as well as the variability of colour contrast (e.g. *Col.kurtosis, VCA.CVS*) and a decrease of the average colour contrast (e.g. *Col.mean*) as well as colour pattern regularity (e.g. *CAA.Qc*).

We found no differences in factor values of backgrounds between groups with different levels of chemical defences [difference (+-95% CI)] (Fig. 2c, Table S7) at either 2cm or 30cm. However, all groups showed increased factor values at 2cm compared to 30cm (Fig 2c; undefended: (0.50 (0.07 / 0.91)); toxic moderate unpalatable: toxic highly unpalatable species (0.34 (0.08 / 0.59)). No difference in the variability of factor values was detected between the background of groups at either viewing distance, and background variability was the same at 2cm and 30cm for all groups (Table S8).

### <u>Factor 4:</u> *Increases in achromatic patch contrast correlate with decreases in achromatic boundary contrast, but increases in chromatic boundary contrast variability*

Factor 4 (Fig. 3d) describes a correlation between achromatic average patch contrast (e.g. *VCA.ML*) and measures of achromatic boundary contrast (e.g. *Lum.mean, BSA.BML*). Specifically, increases in mean luminance contrast relative to the size of colour pattern elements coincide with a decrease in the average luminance contrast of boundaries between colour pattern elements. Furthermore, decreases in the mean luminance contrast of pattern boundaries correlate with an increase in chromatic boundary contrast variability relative to the mean (e.g*. BSA.BCVSsat*).

We did not find any differences in factor values between the backgrounds of species with different levels of chemical defences at either 2cm, or 30cm [difference (+-95% CI)] (Fig. 2d, Table S9). However, while undefended species showed no difference in factor values of backgrounds between viewing distances (0.21 (−0.25 / 0.69)), both toxic and medium unpalatable species (0.53 (0.25 / 0.81)), as well as toxic and highly unpalatable species (0.42 (0.16 / 0.70)), had higher factor values for their visual backgrounds at 2cm compared to 30cm. No differences in background variability between groups were detected at either viewing distance (Table S10), except for undefended species having more variable backgrounds at 30cm compared to toxic and moderately defended species (0.45 (0.07/ 0.90). The variability of visual backgrounds did not differ between viewing distances for either group (Table S10).

## Discussion

Our analysis captured four latent variables describing correlations between the spatiochromatic properties of visual backgrounds on which animals from 12 species of nudibranch molluscs were found (Fig. 2). In agreement with one of our predictions derived from the ‘escape and radiate’ hypothesis, these latent variables showed that nudibranch species with chemical defences, irrespective of their relative strength, were found on visual backgrounds distinct in their appearance from backgrounds of undefended species according to the physiological limitations of a potential predator, a triggerfish (*Rhinecanthus aculeatus*) (Fig. 2a). However, while some visual properties of backgrounds varied significantly more up close (2cm) than from further away (30cm), we did not find any differences in the variability of visual backgrounds between species with different levels of chemical defences. This lack of among-species variability indicates that chemical defences and their relative strength do not correspond to increased among-species background variability in our dataset, contrary to the predictions derived from the ‘escape and radiate’ hypothesis (Merilaita & Tullberg, 2005; Thompson, 1989). Instead, we suggest that chemical defences in Dorid nudibranchs coincide with broad, yet equally variable, differences in visual background habitats.

As shown by the composition of each factor (Fig. 2), multicomponent descriptors of complex phenotypes can be difficult to reduce to a low-dimensional, intuitive, spatiochromatic property that adequately captures the underlying complexity and links it to perceptual and functional properties of colouration. Unlike numerical classifiers from artificial neuronal networks or other machine learning approaches used in high-dimensional image analyses for computer vision (e.g. Bileschi et al., 2007; Talas et al., 2020)), each parameter in our colour pattern space describes a specific spatiochromatic property (van den Berg, Troscianko, et al., 2020). Therefore, despite a range of complexity in associations with response variables (see, e.g. van den Berg et al., 2022) for discussion), we can make assumptions about the perceptual processes associated with our latent predictors.

Visual backgrounds of chemically defended species in our study can be broadly characterised by the presence of increased colour and luminance contrast between objects and surfaces when compared to visual backgrounds of undefended species (Fig. 2a). This difference in the visual appearance of background habitats to a potential predator was equally as strong when viewed from 2cm than at 30cm. Therefore, backgrounds of chemically defended species remained more spatiochromatically contrasting than those of undefended species (Fig. 2a), even at distances where substantial amounts of spatial information would be lost to a potential predator. This persistence of spatiochromatic variability in visual backgrounds across viewing distances is of interest in the context of distance-dependent selection pressures shaping the ecology and evolution of colour pattern diversity in prey communities (e.g. Barnett et al., 2016; Endler, 1978; van den Berg et al., 2022; van den Berg, Endler, et al., 2023). For example, imperfect mimicry among- and colour pattern polymorphism within aposematic species could be shaped by their adaptive value in the context of predator perception at multiple viewing distances. Persisting variability of visual backgrounds across viewing distances could be selecting for perceptual similarities between animals and backgrounds in a more general way than previously considered.

Aposematism is assumed to be widespread in Dorid nudibranchs (Rudman, 1991), with bold colours and patterns coinciding with the presence of chemical defences (Cortesi & Cheney, 2010; Winters et al., 2022; Winters, Wilson, et al., 2018). However, the coincidence of secondary defences and boldly contrasting animal colouration (e.g. van den Berg, Endler, et al., 2023) does not mean that the colours and patterns displayed by a species necessarily serve a warning function (see Summers et al., 2015; White & Umbers, 2021) for review). Instead, the function of bold colouration in an environment with increased colour and luminance contrast is likely complex and might even assist in camouflage (Endler, 1978; Marshall & Stevens, 2014; van den Berg, Endler, et al., 2023). The detectability of an aposematic animal is determined by its appearance against its visual background and is likely fine-tuned to the cost-benefit trade-offs of increased predator encounters (e.g. Barnett et al., 2016; van den Berg, Endler, et al., 2023). Therefore, assuming substantial (yet variable) degrees of signalling honesty in Dorid nudibranchs (e.g. Cortesi & Cheney, 2010), our study suggests broadly generalisable correlations between the presence of chemical defences in aposematic species and the spatiochromatic properties of their visual habitat. However, these constraints can be variably explained by the need for efficient camouflage (e.g. Barnett et al., 2016; Speed, Ruxton, et al., 2010) or signalling efficacy (e.g. Endler & Mappes, 2004; Speed, Brockhurst, et al., 2010) at variable viewing distances. How specific spatiochromatic properties of visual backgrounds highlighted by our study impact the signalling function of aposematic colouration and camouflage in the considered species would be of great interest in future behavioural experimentation studies.

Nudibranchs mainly use chemotaxis to move around their environment to find food and mates, with visual input only relevant for basic phototaxis, such as determining daytime or detecting shelter (Eakin et al., 1967). Therefore, unlike other aposematic species, such as insects or frogs (Higginson et al., 2012; Rojas, 2017), nudibranchs will unlikely choose resting and foraging microhabitats based on visual cues. Therefore, it is likely that the backgrounds on which species are found are the indirect rather than direct consequence of correlations between their habitat’s visual appearance and other sensory modes and selective pressures shaping the efficacy of visual defences. Thus, finding chemically defended species on distinct visual backgrounds fits well with assumed radiation in feeding ecology, enabling the acquisition of secondary defences (e.g. Winters, White, et al., 2018).

To our knowledge, no study has quantified the mobility and the diets of many nudibranch species (but see Rudman & Bergquist, 2007), and the spatiotemporal distribution of food sources remains poorly known. Some of the most unpalatable nudibranchs in this study are known to forage on sponges containing highly potent secondary metabolites (e.g. latrunculin a) at least at some stage during their lifetime (e.g. *Chromodoris elisabethina*) (Cheney et al., 2016). In contrast, others can synthesise compounds *de novo* (Cimino & Ghiselin, 1999). Nudibranchs with more specialist diets may need to increase mobility to find uncommon food sources. The sponge diet consequently allows the animals to maintain functional levels of secondary metabolites or pigments required to maintain salient aposematic colour patterns. Background specialisation might, therefore, be increased in species dependent on such supplemental food sources and could influence within and among-species background variability. However, whether the presence of secondary defences in the context of visual background habitats also coincides with changes in defensive animal colouration remains an exciting and crucial avenue for future research.

The mechanisms underlying the ‘escape and radiate’ hypothesis are often unclear, with little empirical evidence of the process (e.g. Suchan & Alvarez, 2015). For example, evidence of increased background variability could only be present during certain stages of (rapid) speciation in aposematic animals. More extensive sampling of primary and secondary defences and corresponding visual backgrounds within and among species, considering each species’ geographic distribution, would be of great interest for future research. Combined with an increased resolution of existing heterobranch phylogenies (e.g. Layton et al., 2018, 2020), this would enable a more detailed investigation into the presence and scale of animal background and colour pattern diversification and its correlation with secondary defences.

The direction, speed and extent by which natural selection would constrain correlations between the strength of secondary defences and background habitat is likely determined by multiple, currently poorly understood factors such as the fitness benefits of specific colour patterns across different visual backgrounds, the diversity of visual backgrounds inhabited by a given species, the protective value of secondary compounds across complex predator communities and the heritability of visual phenotypes, to name a few. Nevertheless, our study provides empirical support to the possibility of secondary defences coinciding with generalisable differences in visual background habitats in a complex community of nudibranch molluscs. We demonstrate this by employing sophisticated methodology aimed at reflecting visual processing of ecologically relevant observers and utilising a high-dimensional, highly differentiated approach to quantifying spatiochromatic properties of visual backgrounds.

## Acknowledgements

We thank various volunteers for assistance with fieldwork and image analysis. We are grateful for the High-Performance Computing (HPC) infrastructure at The University of Queensland (Wiener & Awoonga) and Dr. Simone Blomberg, who provided infrastructure that contributed to the computing of image statistics.

## Data availability

The data is available on UQ’s digital repository eSpace: https://doi.org/10.48610/8f7b70e

## Author contributions

CPvdB conceived the original study with KLC & JAE providing critical input; CPvdB collected and analysed the data; CPvdB and KLC led the writing of the manuscript. All authors contributed critically to the drafts and gave final approval for publication.

---

**Figure S1.**
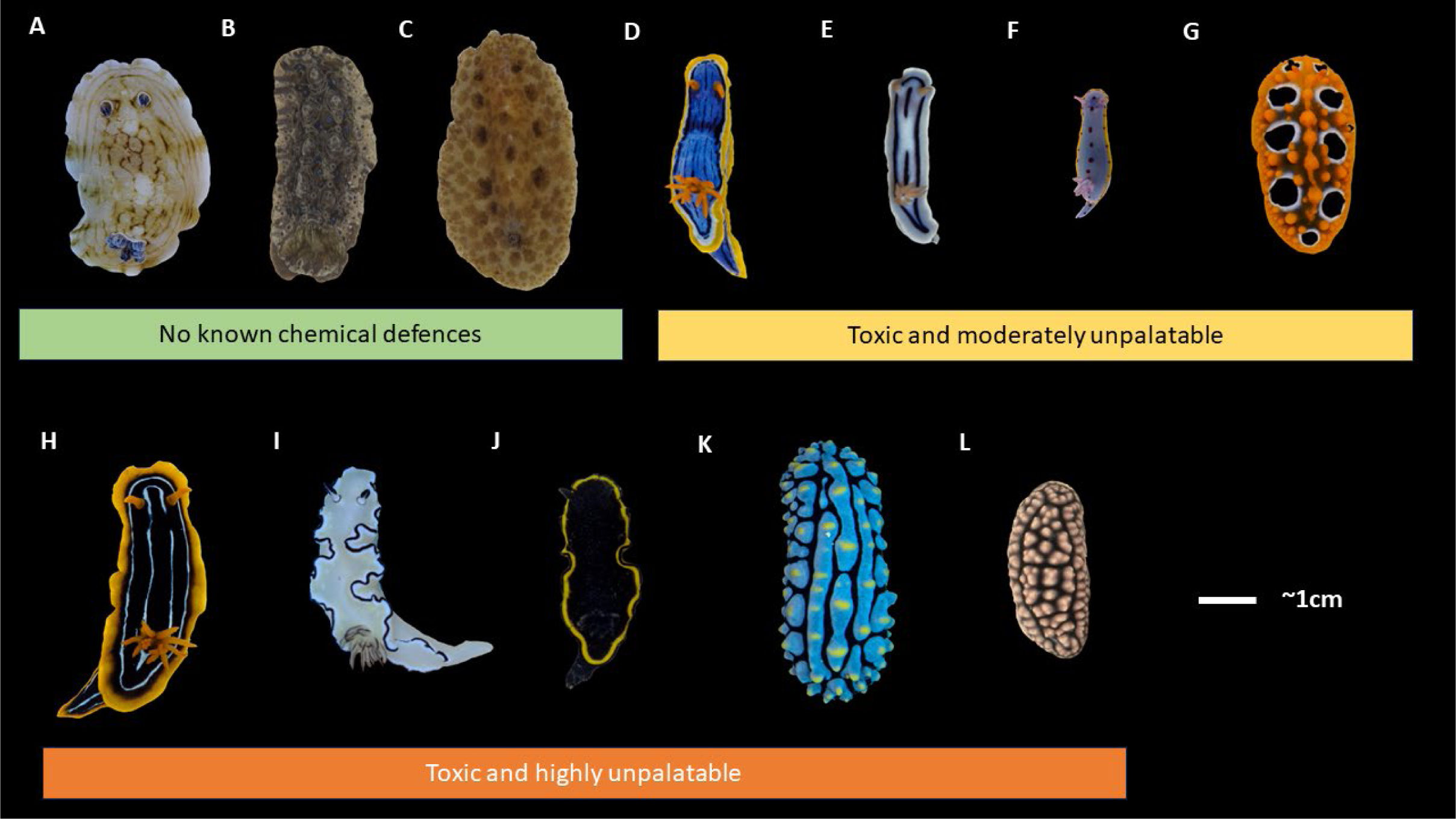
Representative photographs of the 12 species used in this study grouped into categories of chemical defences based on whole-body extract assays with palaemon shrimp to assess unpalatability (1-Effective Dose, ED_50_) and brine shrimp to assess toxicity (1-Lethal Dose, LD_50_) values: A) *Aphelodoris varia;* B) *Dendrodoris krusensterni*; C) *Discodoris sp*; D) *Chromodoris elisabethina*; E) *Chromodoris lochi;* F) *Hypselodoris bennetti;* G) *Phyllidia ocellata*; H) *Chromodoris kuiteri*; I) *Doriprismatica atromarginata;* J) *Glossodoris vespa*; K) *Phyllidia varicosa;* L) *Phyllidiella pustulosa*.

**Figure S2.**
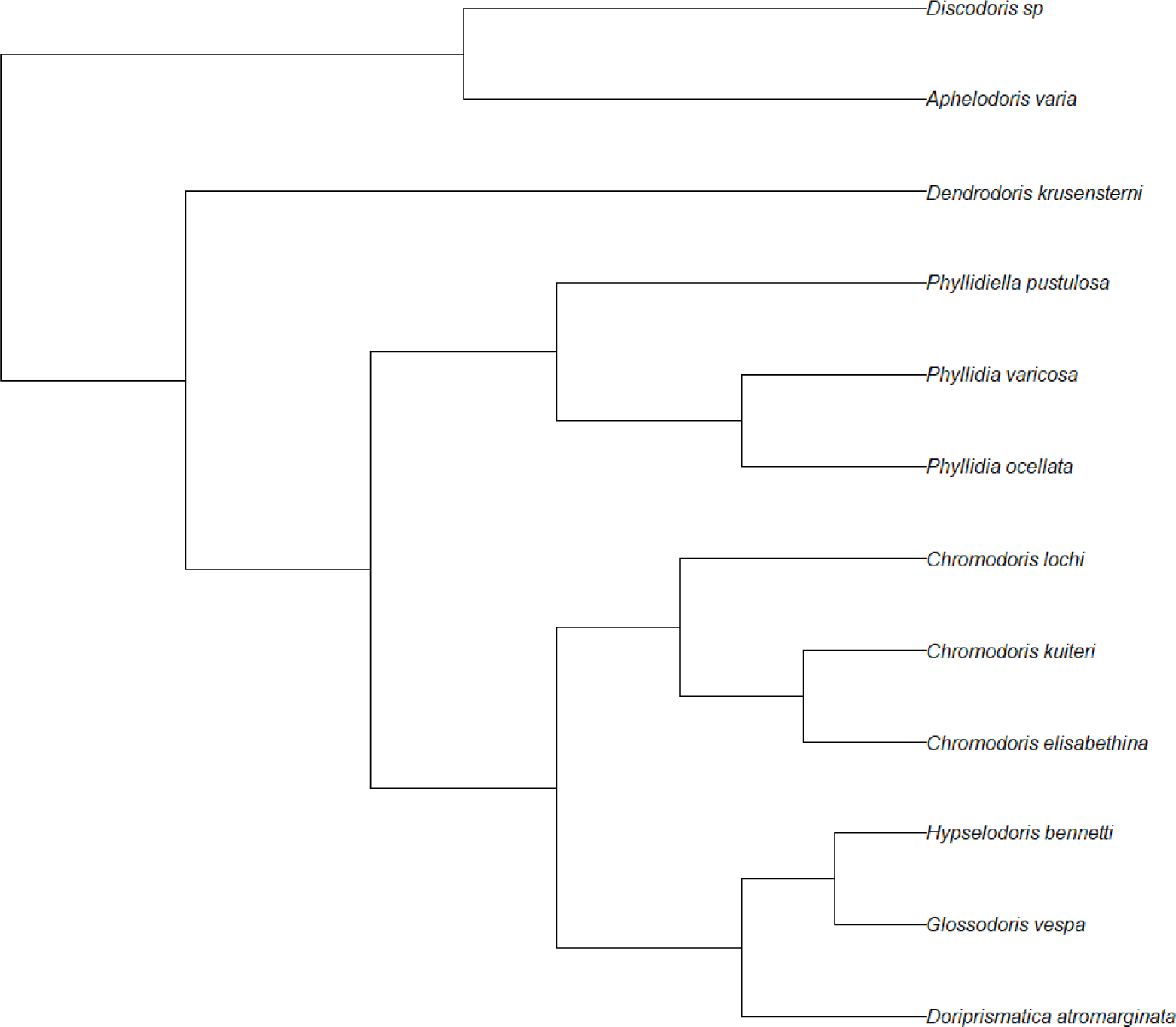
Phylogenetic tree used in this study, modified from Cheney et al. 2014 as described in ‘Methods’.

**Table S1.** Summary table of all individuals in the dataset used for this study

| Species | Unpalatability (1-ED<sub>50</sub>) | Toxicity (1-LD<sub>50</sub>) | Sampling site |  |  |  |
| --- | --- | --- | --- | --- | --- | --- |
|  |  |  | Sunshine Coast, QLD | Gold Coast, QLD | Nelson Bay, NSW | Grand Total |
| *Aphelodoris varia* | 0 | 0 | 0 | 0 | 22 | 22 |
| *Chromodoris elisabethina* | 0.55 | 1 | 21 | 0 | 0 | 21 |
| *Chromodoris kuiteri* | 0.74 | 1 | 10 | 7 | 0 | 17 |
| *Chromodoris lochi* | 0.61 | 1 | 3 | 0 | 0 | 3 |
| *Dendrodoris krusensterni* | 0 | 0 | 0 | 2 | 5 | 7 |
| *Discodoris sp.* | 0 | 0 | 0 | 0 | 13 | 13 |
| *Doriprismatica atromarginata* | 0.74 | 0.93 | 16 | 0 | 11 | 27 |
| *Glossodoris vespa* | 0.87 | 0.29 | 15 | 0 | 0 | 15 |
| *Hypselodoris bennetti* | 0.73 | 0.88 | 0 | 0 | 10 | 10 |
| *Phyllidia ocellata* | 0.63 | 0.4 | 23 | 0 | 0 | 23 |
| *Phyllidia varicosa* | 0.92 | 0.53 | 8 | 0 | 0 | 8 |
| *Phyllidiella pustulosa* | 0.86 | 0.14 | 18 | 0 | 0 | 18 |

**Table S2.**
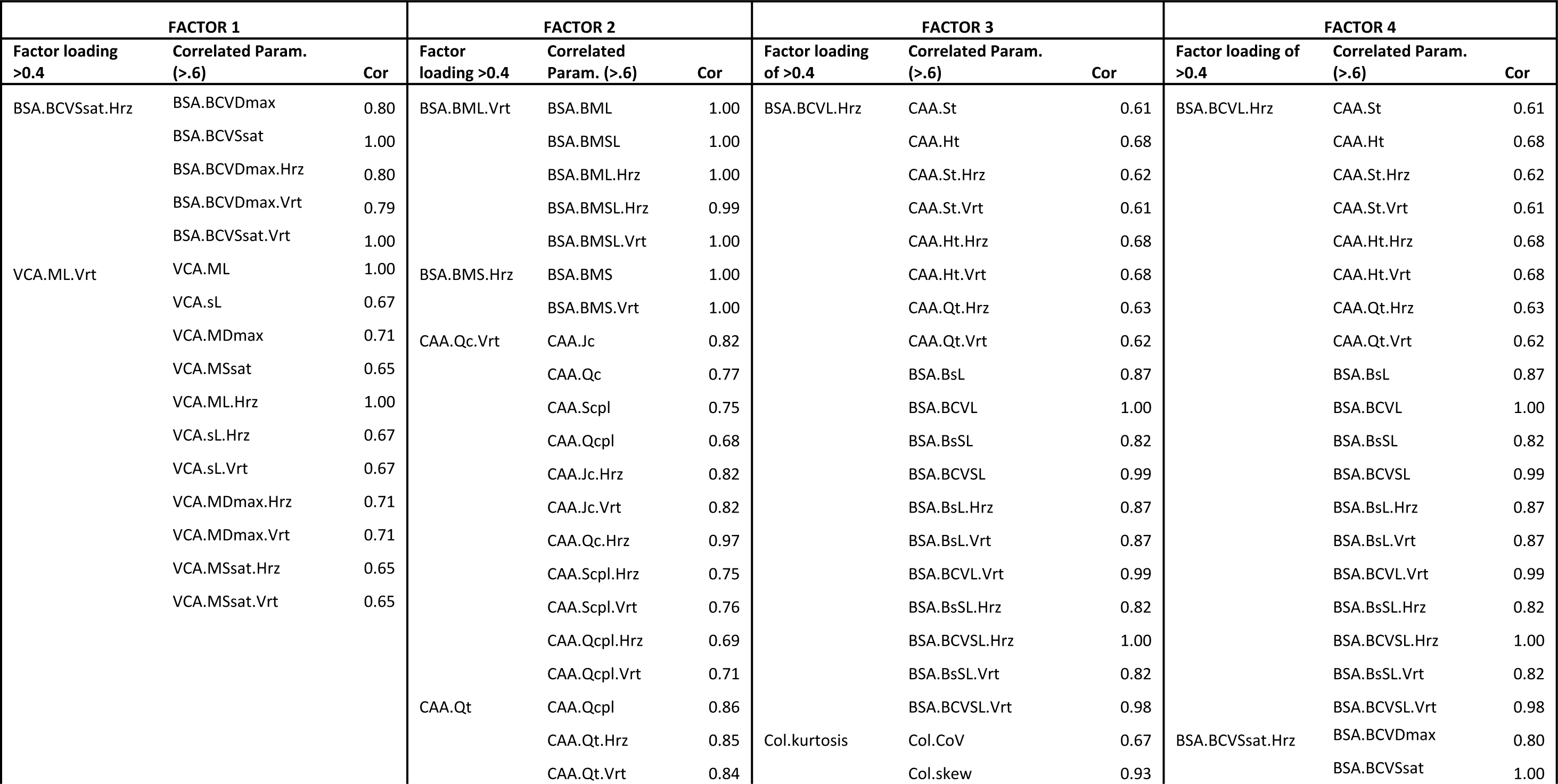
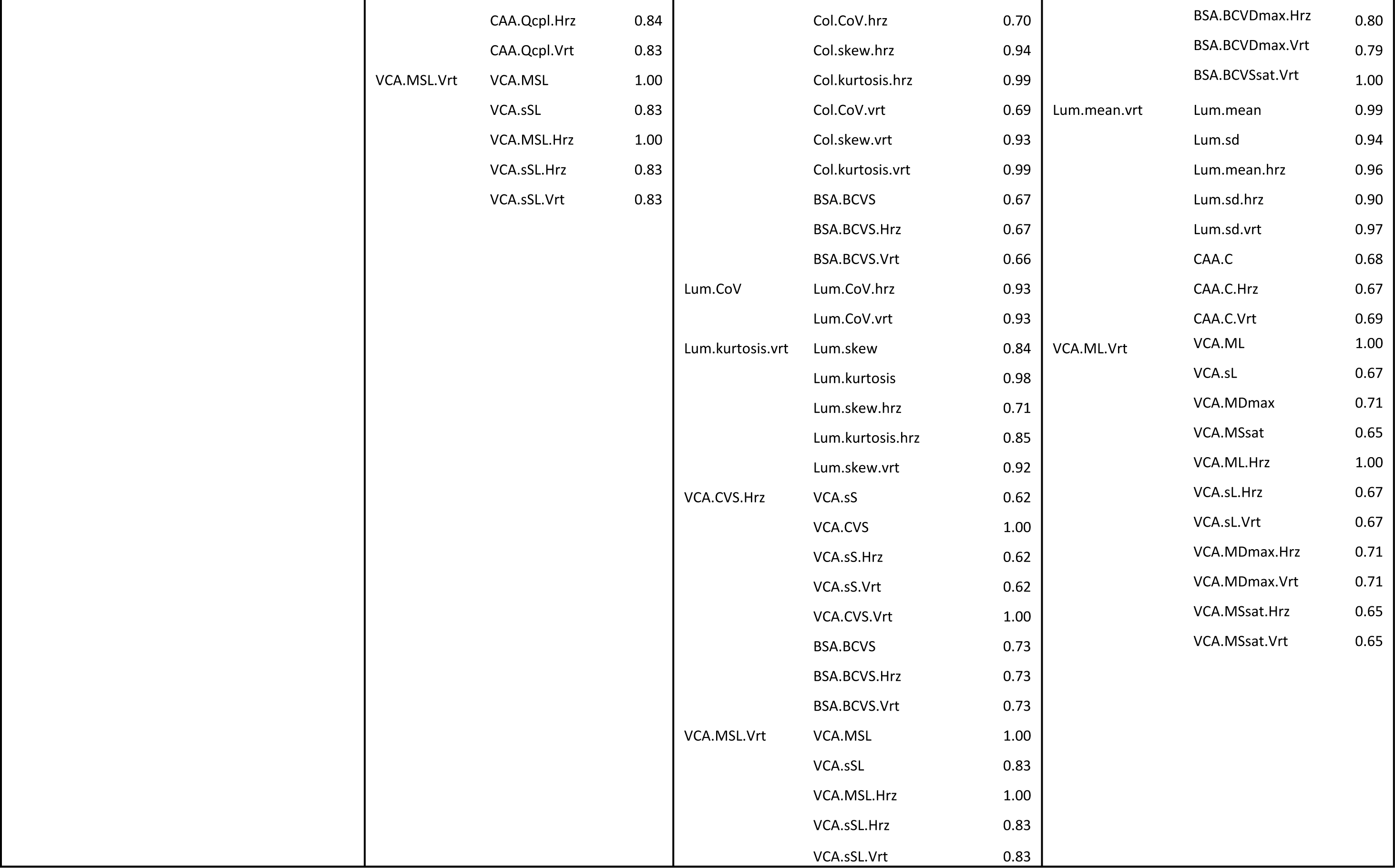
Summary table of all >.6 Pearson correlation values between parameters with >0.4 loadings for each factor.

**Table S3.** Pairwise contrasts for the model investigating latent factor 1 expressed as the median differences between groups with different strengths of chemical defences (see ‘Methods’ for details). The effect size of pairwise differences increases with increasing deviation of such differences from zero, and the robustness of the result increases with decreasing degree of overlap of the 95% Credible Intervals (CIs) with zero.

| **Factor 1: Contrasts** | **Difference** | **Lower CI** | **Upper CI** |
| --- | --- | --- | --- |
| Undefended at 2cm - Toxic and moderately unpalatable at 2cm | -1.53 | -2.28 | -0.78 |
| Undefended at 2cm - Toxic and highly unpalatable at 2cm | -1.44 | -2.18 | -0.73 |
| Undefended at 2cm - Undefended at 30cm | -0.18 | -0.61 | 0.27 |
| Undefended at 2cm - Toxic and moderately unpalatable at 30cm | -1.23 | -1.99 | -0.46 |
| Undefended at 2cm - Toxic and highly unpalatable at 30cm | -1.35 | -2.10 | -0.65 |
| Toxic and moderately unpalatable at 2cm - Toxic and highly unpalatable at 2cm | 0.08 | -0.38 | 0.54 |
| Toxic and moderately unpalatable at 2cm - Undefended at 30cm | 1.35 | 0.58 | 2.15 |
| Toxic and moderately unpalatable at 2cm - Toxic and moderately unpalatable at 30cm | 0.30 | -0.02 | 0.63 |
| Toxic and moderately unpalatable at 2cm - Toxic and highly unpalatable at 30cm | 0.17 | -0.28 | 0.65 |
| Toxic and highly unpalatable at 2cm - Undefended at 30cm | 1.26 | 0.47 | 2.04 |
| Toxic and highly unpalatable at 2cm - Toxic and moderately unpalatable at 30cm | 0.22 | -0.29 | 0.70 |
| Toxic and highly unpalatable at 2cm - Toxic and highly unpalatable at 30cm | 0.09 | -0.19 | 0.38 |
| Undefended at 30cm - Toxic and moderately unpalatable at 30cm | -1.05 | -1.83 | -0.23 |
| Undefended at 30cm - Toxic and highly unpalatable at 30cm | -1.17 | -1.92 | -0.37 |
| Toxic and moderately unpalatable at 30cm - Toxic and highly unpalatable at 30cm | -0.13 | -0.63 | 0.36 |

**Table S4.** Pairwise variance contrasts for the model investigating latent factor 1 expressed as the median differences of the residual standard deviation on the original scale (back-transformed from the log scale) between groups with different strengths of chemical defences (see ‘Methods’ for details). The effect size of pairwise differences increases with increasing deviation of such differences from zero, and the robustness of the result increases with decreasing degree of overlap of the 95% Credible Intervals (CIs) with zero.

| **Factor 1: variance contrasts** | **Difference** | **Lower CI** | **Upper CI** |
| --- | --- | --- | --- |
| Undefended at 2cm - Toxic and moderately unpalatable at 2cm | 0.21 | -0.07 | 0.55 |
| Undefended at 2cm - Toxic and highly unpalatable at 2cm | -0.03 | -0.30 | 0.31 |
| Undefended at 2cm - Undefended at 30cm | -0.12 | -0.53 | 0.22 |
| Undefended at 2cm - Toxic and moderately unpalatable at 30cm | -0.21 | -0.67 | 0.12 |
| Undefended at 2cm - Toxic and highly unpalatable at 30cm | 0.13 | -0.09 | 0.33 |
| Toxic and moderately unpalatable at 2cm - Toxic and highly unpalatable at 2cm | -0.23 | -0.49 | 0.01 |
| Toxic and moderately unpalatable at 2cm - Undefended at 30cm | -0.33 | -0.76 | 0.01 |
| Toxic and moderately unpalatable at 2cm - Toxic and moderately unpalatable at 30cm | -0.42 | -1.09 | 0.08 |
| Toxic and moderately unpalatable at 2cm - Toxic and highly unpalatable at 30cm | -0.08 | -0.45 | 0.22 |
| Toxic and highly unpalatable at 2cm - Undefended at 30cm | -0.09 | -0.52 | 0.25 |
| Toxic and highly unpalatable at 2cm - Toxic and moderately unpalatable at 30cm | -0.19 | -0.81 | 0.26 |
| Toxic and highly unpalatable at 2cm - Toxic and highly unpalatable at 30cm | 0.16 | -0.26 | 0.51 |
| Undefended at 30cm - Toxic and moderately unpalatable at 30cm | -0.09 | -0.71 | 0.44 |
| Undefended at 30cm - Toxic and highly unpalatable at 30cm | 0.25 | -0.15 | 0.69 |
| Toxic and moderately unpalatable at 30cm - Toxic and highly unpalatable at 30cm | 0.34 | -0.06 | 0.85 |

**Table S5.** Pairwise contrasts for the model investigating latent factor 2 expressed as the median differences between groups with different strengths of chemical defences (see ‘Methods’ for details). The effect size of pairwise differences increases with increasing deviation of such differences from zero, and the robustness of the result increases with decreasing degree of overlap of the 95% Credible Intervals (CIs) with zero.

| **Factor 2: Contrasts** | **Difference** | **Lower CI** | **Upper CI** |
| --- | --- | --- | --- |
| Undefended at 2cm - Toxic and moderately unpalatable at 2cm | -0.1226 | -0.614 | 0.402 |
| Undefended at 2cm - Toxic and highly unpalatable at 2cm | -0.1056 | -0.617 | 0.394 |
| Undefended at 2cm - Undefended at 30cm | 1.0072 | 0.568 | 1.482 |
| Undefended at 2cm - Toxic and moderately unpalatable at 30cm | 0.8871 | 0.352 | 1.417 |
| Undefended at 2cm - Toxic and highly unpalatable at 30cm | 0.6930 | 0.125 | 1.194 |
| Toxic and moderately unpalatable at 2cm - Toxic and highly unpalatable at 2cm | 0.0175 | -0.333 | 0.347 |
| Toxic and moderately unpalatable at 2cm - Undefended at 30cm | 1.1337 | 0.552 | 1.705 |
| Toxic and moderately unpalatable at 2cm - Toxic and moderately unpalatable at 30cm | 1.0052 | 0.668 | 1.312 |
| Toxic and moderately unpalatable at 2cm - Toxic and highly unpalatable at 30cm | 0.8158 | 0.410 | 1.164 |
| Toxic and highly unpalatable at 2cm - Undefended at 30cm | 1.1151 | 0.552 | 1.686 |
| Toxic and highly unpalatable at 2cm - Toxic and moderately unpalatable at 30cm | 0.9896 | 0.615 | 1.363 |
| Toxic and highly unpalatable at 2cm - Toxic and highly unpalatable at 30cm | 0.7997 | 0.473 | 1.105 |
| Undefended at 30cm - Toxic and moderately unpalatable at 30cm | -0.1239 | -0.709 | 0.462 |
| Undefended at 30cm - Toxic and highly unpalatable at 30cm | -0.3172 | -0.971 | 0.230 |
| Toxic and moderately unpalatable at 30cm - Toxic and highly unpalatable at 30cm | -0.1898 | -0.615 | 0.215 |

**Table S6.**
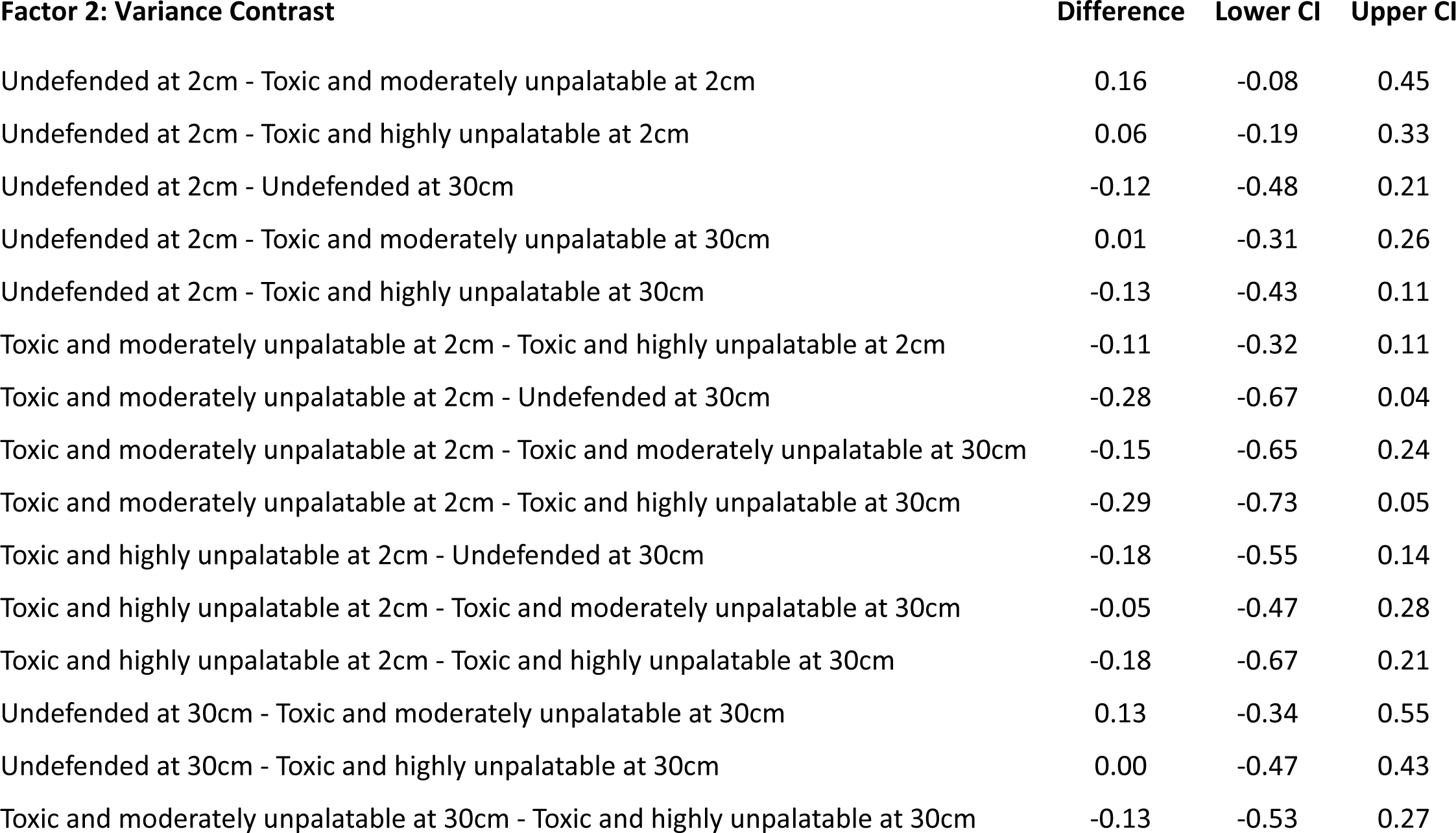
Pairwise variance contrasts for the model investigating latent factor 2 expressed as the median differences of the residual standard deviation on the original scale (back-transformed from the log scale) between groups with different strengths of chemical defences (see ‘Methods’ for details). The effect size of pairwise differences increases with increasing deviation of such differences from zero, and the robustness of the result increases with decreasing degree of overlap of the 95% Credible Intervals (CIs) with zero.

| **Factor 2: Variance Contrast** | **Difference** | **Lower CI** | **Upper CI** |
| --- | --- | --- | --- |
| Undefended at 2cm - Toxic and moderately unpalatable at 2cm | 0.16 | -0.08 | 0.45 |
| Undefended at 2cm - Toxic and highly unpalatable at 2cm | 0.06 | -0.19 | 0.33 |
| Undefended at 2cm - Undefended at 30cm | -0.12 | -0.48 | 0.21 |
| Undefended at 2cm - Toxic and moderately unpalatable at 30cm | 0.01 | -0.31 | 0.26 |
| Undefended at 2cm - Toxic and highly unpalatable at 30cm | -0.13 | -0.43 | 0.11 |
| Toxic and moderately unpalatable at 2cm - Toxic and highly unpalatable at 2cm | -0.11 | -0.32 | 0.11 |
| Toxic and moderately unpalatable at 2cm - Undefended at 30cm | -0.28 | -0.67 | 0.04 |
| Toxic and moderately unpalatable at 2cm - Toxic and moderately unpalatable at 30cm | -0.15 | -0.65 | 0.24 |
| Toxic and moderately unpalatable at 2cm - Toxic and highly unpalatable at 30cm | -0.29 | -0.73 | 0.05 |
| Toxic and highly unpalatable at 2cm - Undefended at 30cm | -0.18 | -0.55 | 0.14 |
| Toxic and highly unpalatable at 2cm - Toxic and moderately unpalatable at 30cm | -0.05 | -0.47 | 0.28 |
| Toxic and highly unpalatable at 2cm - Toxic and highly unpalatable at 30cm | -0.18 | -0.67 | 0.21 |
| Undefended at 30cm - Toxic and moderately unpalatable at 30cm | 0.13 | -0.34 | 0.55 |
| Undefended at 30cm - Toxic and highly unpalatable at 30cm | 0.00 | -0.47 | 0.43 |
| Toxic and moderately unpalatable at 30cm - Toxic and highly unpalatable at 30cm | -0.13 | -0.53 | 0.27 |

**Table S7.** Pairwise contrasts for the model investigating latent factor 3 expressed as the median differences between groups with different strengths of chemical defences (see ‘Methods’ for details). The effect size of pairwise differences increases with increasing deviation of such differences from zero, and the robustness of the result increases with decreasing degree of overlap of the 95% Credible Intervals (CIs) with zero.

| **Factor 3: Contrasts** | **Difference** | **Lower CI** | **Upper CI** |
| --- | --- | --- | --- |
| Undefended at 2cm - Toxic and moderately unpalatable at 2cm | 0.23 | -0.27 | 0.69 |
| Undefended at 2cm - Toxic and highly unpalatable at 2cm | 0.18 | -0.26 | 0.66 |
| Undefended at 2cm - Undefended at 30cm | 0.50 | 0.07 | 0.91 |
| Undefended at 2cm - Toxic and moderately unpalatable at 30cm | 0.50 | 0.00 | 1.01 |
| Undefended at 2cm - Toxic and highly unpalatable at 30cm | 0.51 | 0.02 | 0.97 |
| Toxic and moderately unpalatable at 2cm - Toxic and highly unpalatable at 2cm | -0.05 | -0.37 | 0.24 |
| Toxic and moderately unpalatable at 2cm - Undefended at 30cm | 0.26 | -0.31 | 0.82 |
| Toxic and moderately unpalatable at 2cm - Toxic and moderately unpalatable at 30cm | 0.27 | -0.03 | 0.56 |
| Toxic and moderately unpalatable at 2cm - Toxic and highly unpalatable at 30cm | 0.28 | -0.08 | 0.60 |
| Toxic and highly unpalatable at 2cm - Undefended at 30cm | 0.32 | -0.24 | 0.84 |
| Toxic and highly unpalatable at 2cm - Toxic and moderately unpalatable at 30cm | 0.32 | -0.02 | 0.69 |
| Toxic and highly unpalatable at 2cm - Toxic and highly unpalatable at 30cm | 0.33 | 0.07 | 0.56 |
| Undefended at 30cm - Toxic and moderately unpalatable at 30cm | 0.01 | -0.59 | 0.60 |
| Undefended at 30cm - Toxic and highly unpalatable at 30cm | 0.01 | -0.53 | 0.60 |
| Toxic and moderately unpalatable at 30cm - Toxic and highly unpalatable at 30cm | 0.01 | -0.35 | 0.40 |

**Table S8.** Pairwise variance contrasts for the model investigating latent factor 3 expressed as the median differences of the residual standard deviation on the original scale (back-transformed from the log scale) between groups with different strengths of chemical defences (see ‘Methods’ for details). The effect size of pairwise differences increases with increasing deviation of such differences from zero, and the robustness of the result increases with decreasing degree of overlap of the 95% Credible Intervals (CIs) with zero.

| **Factor 3: Variance Contrast** | **Estimate** | **Lower CI** | **Upper CI** |
| --- | --- | --- | --- |
| Undefended at 2cm - Toxic and moderately unpalatable at 2cm | 0.15 | -0.10 | 0.44 |
| Undefended at 2cm - Toxic and highly unpalatable at 2cm | 0.09 | -0.15 | 0.38 |
| Undefended at 2cm - Undefended at 30cm | -0.19 | -0.62 | 0.15 |
| Undefended at 2cm - Toxic and moderately unpalatable at 30cm | -0.12 | -0.46 | 0.14 |
| Undefended at 2cm - Toxic and highly unpalatable at 30cm | 0.00 | -0.24 | 0.19 |
| Toxic and moderately unpalatable at 2cm - Toxic and highly unpalatable at 2cm | -0.06 | -0.26 | 0.14 |
| Toxic and moderately unpalatable at 2cm - Undefended at 30cm | -0.34 | -0.78 | -0.01 |
| Toxic and moderately unpalatable at 2cm - Toxic and moderately unpalatable at 30cm | -0.27 | -0.78 | 0.15 |
| Toxic and moderately unpalatable at 2cm - Toxic and highly unpalatable at 30cm | -0.15 | -0.55 | 0.15 |
| Toxic and highly unpalatable at 2cm - Undefended at 30cm | -0.28 | -0.72 | 0.05 |
| Toxic and highly unpalatable at 2cm - Toxic and moderately unpalatable at 30cm | -0.21 | -0.67 | 0.15 |
| Toxic and highly unpalatable at 2cm - Toxic and highly unpalatable at 30cm | -0.09 | -0.53 | 0.25 |
| Undefended at 30cm - Toxic and moderately unpalatable at 30cm | 0.08 | -0.43 | 0.58 |
| Undefended at 30cm - Toxic and highly unpalatable at 30cm | 0.19 | -0.23 | 0.66 |
| Toxic and moderately unpalatable at 30cm - Toxic and highly unpalatable at 30cm | 0.11 | -0.23 | 0.51 |

**Table S9.** Pairwise contrasts for the model investigating latent factor 4 expressed as the median differences between groups with different strengths of chemical defences (see ‘Methods’ for details). The effect size of pairwise differences increases with increasing deviation of such differences from zero, and the robustness of the result increases with decreasing degree of overlap of the 95% Credible Intervals (CIs) with zero.

| **Factor 4: Contrasts** | **Estimate** | **Lower CI** | **Upper CI** |
| --- | --- | --- | --- |
| Undefended at 2cm - Toxic and moderately unpalatable at 2cm | 0.21 | -0.51 | 0.96 |
| Undefended at 2cm - Toxic and highly unpalatable at 2cm | 0.31 | -0.38 | 1.04 |
| Undefended at 2cm - Undefended at 30cm | 0.21 | -0.25 | 0.69 |
| Undefended at 2cm - Toxic and moderately unpalatable at 30cm | 0.75 | 0.02 | 1.48 |
| Undefended at 2cm - Toxic and highly unpalatable at 30cm | 0.73 | 0.05 | 1.51 |
| Toxic and moderately unpalatable at 2cm - Toxic and highly unpalatable at 2cm | 0.09 | -0.31 | 0.53 |
| Toxic and moderately unpalatable at 2cm - Undefended at 30cm | -0.01 | -0.81 | 0.78 |
| Toxic and moderately unpalatable at 2cm - Toxic and moderately unpalatable at 30cm | 0.53 | 0.25 | 0.81 |
| Toxic and moderately unpalatable at 2cm - Toxic and highly unpalatable at 30cm | 0.52 | 0.08 | 0.98 |
| Toxic and highly unpalatable at 2cm - Undefended at 30cm | -0.10 | -0.92 | 0.65 |
| Toxic and highly unpalatable at 2cm - Toxic and moderately unpalatable at 30cm | 0.44 | 0.02 | 0.84 |
| Toxic and highly unpalatable at 2cm - Toxic and highly unpalatable at 30cm | 0.42 | 0.16 | 0.70 |
| Undefended at 30cm - Toxic and moderately unpalatable at 30cm | 0.54 | -0.26 | 1.35 |
| Undefended at 30cm - Toxic and highly unpalatable at 30cm | 0.53 | -0.27 | 1.32 |
| Toxic and moderately unpalatable at 30cm - Toxic and highly unpalatable at 30cm | -0.02 | -0.42 | 0.44 |

**Table S10.** Pairwise variance contrasts for the model investigating latent factor 4 expressed as the median differences of the residual standard deviation on the original scale (back-transformed from the log scale) between groups with different strengths of chemical defences (see ‘Methods’ for details). The effect size of pairwise differences increases with increasing deviation of such differences from zero, and the robustness of the result increases with decreasing degree of overlap of the 95% Credible Intervals (CIs) with zero.

| **Factor 4: Variance Contrasts** | **Estimate** | **Lower CI** | **Upper CI** |
| --- | --- | --- | --- |
| Undefended at 2cm - Toxic and moderately unpalatable at 2cm | 0.08 | -0.19 | 0.40 |
| Undefended at 2cm - Toxic and highly unpalatable at 2cm | 0.07 | -0.19 | 0.39 |
| Undefended at 2cm - Undefended at 30cm | -0.24 | -0.66 | 0.11 |
| Undefended at 2cm - Toxic and moderately unpalatable at 30cm | 0.21 | 0.00 | 0.40 |
| Undefended at 2cm - Toxic and highly unpalatable at 30cm | -0.05 | -0.31 | 0.14 |
| Toxic and moderately unpalatable at 2cm - Toxic and highly unpalatable at 2cm | -0.01 | -0.24 | 0.23 |
| Toxic and moderately unpalatable at 2cm - Undefended at 30cm | -0.32 | -0.80 | 0.04 |
| Toxic and moderately unpalatable at 2cm - Toxic and moderately unpalatable at 30cm | 0.13 | -0.27 | 0.46 |
| Toxic and moderately unpalatable at 2cm - Toxic and highly unpalatable at 30cm | -0.13 | -0.56 | 0.21 |
| Toxic and highly unpalatable at 2cm - Undefended at 30cm | -0.31 | -0.78 | 0.04 |
| Toxic and highly unpalatable at 2cm - Toxic and moderately unpalatable at 30cm | 0.14 | -0.20 | 0.40 |
| Toxic and highly unpalatable at 2cm - Toxic and highly unpalatable at 30cm | -0.12 | -0.60 | 0.25 |
| Undefended at 30cm - Toxic and moderately unpalatable at 30cm | 0.45 | 0.07 | 0.90 |
| Undefended at 30cm - Toxic and highly unpalatable at 30cm | 0.19 | -0.25 | 0.65 |
| Toxic and moderately unpalatable at 30cm - Toxic and highly unpalatable at 30cm | -0.26 | -0.59 | 0.03 |

**Table S11.** Coefficient estimates of the model investigating the scores for latent factor 1 between species of nudibranchs with different levels of chemical defences (N = 12, R^2 = 0.36). Estimates are based on a Student distribution with an identity link for the mean of the response distribution and a log link for its residual standard deviation (Sigma). The estimate is more likely to be non-zero when the credible intervals do not overlap with zero.

| Coefficient | Mean | M. Error | 95% CIs |  |
| --- | --- | --- | --- | --- |
|  |  |  | Low | High |
| Group-level effects |  |  |  |  |
| Phylogenesis |  |  |  |  |
| Sd [Intercept] | 0.24 | 0.18 | 0.01 | 0.66 |
| Species of nudibranch |  |  |  |  |
| Sd [Intercept: Distance 2 cm] | 0.21 | 0.12 | 0.02 | 0.48 |
| Sd [Distance 30 cm] | 0.12 | 0.10 | 0.00 | 0.36 |
| Sigma: Sd [Intercept: Distance 2 cm] | 0.12 | 0.09 | 0.00 | 0.35 |
| Sigma: Sd [Distance 30 cm] | 0.14 | 0.11 | 0.01 | 0.41 |
| Population-level effects |  |  |  |  |
| Intercept [Undefended, 2 cm] | -1.08 | 0.28 | -1.63 | -0.51 |
| Sigma_Intercept [Undefended, 2 cm] | -0.24 | 0.16 | -0.53 | 0.08 |
| Chemical defence [Toxic and moderately unpalatable] | 1.52 | 0.37 | 0.73 | 2.24 |
| Chemical defence [Toxic and highly unpalatable] | 1.44 | 0.36 | 0.73 | 2.18 |
| Distance [30 cm] | 0.18 | 0.22 | -0.25 | 0.63 |
| Chemical defence [Toxic and moderately unpalatable] x Distance [30 cm] | -0.48 | 0.28 | -1.02 | 0.07 |
| Chemical defence [Toxic and highly unpalatable] x Distance [30 cm] | -0.27 | 0.27 | -0.80 | 0.25 |
| Sigma: Chemical defence [Toxic and moderately unpalatable] | -0.30 | 0.21 | -0.76 | 0.1 |
| Sigma: Chemical defence [Toxic and highly unpalatable] | 0.04 | 0.19 | -0.36 | 0.38 |
| Sigma: Distance [30cm] | 0.13 | 0.21 | -0.27 | 0.55 |
| Sigma: Chemical defence [Toxic and moderately unpalatable] x Distance [30 cm] | 0.09 | 0.28 | -0.47 | 0.65 |
| Sigma: Chemical defence [Toxic and highly unpalatable] x Distance [30 cm] | -0.32 | 0.26 | -0.85 | 0.18 |

**Table S12.**
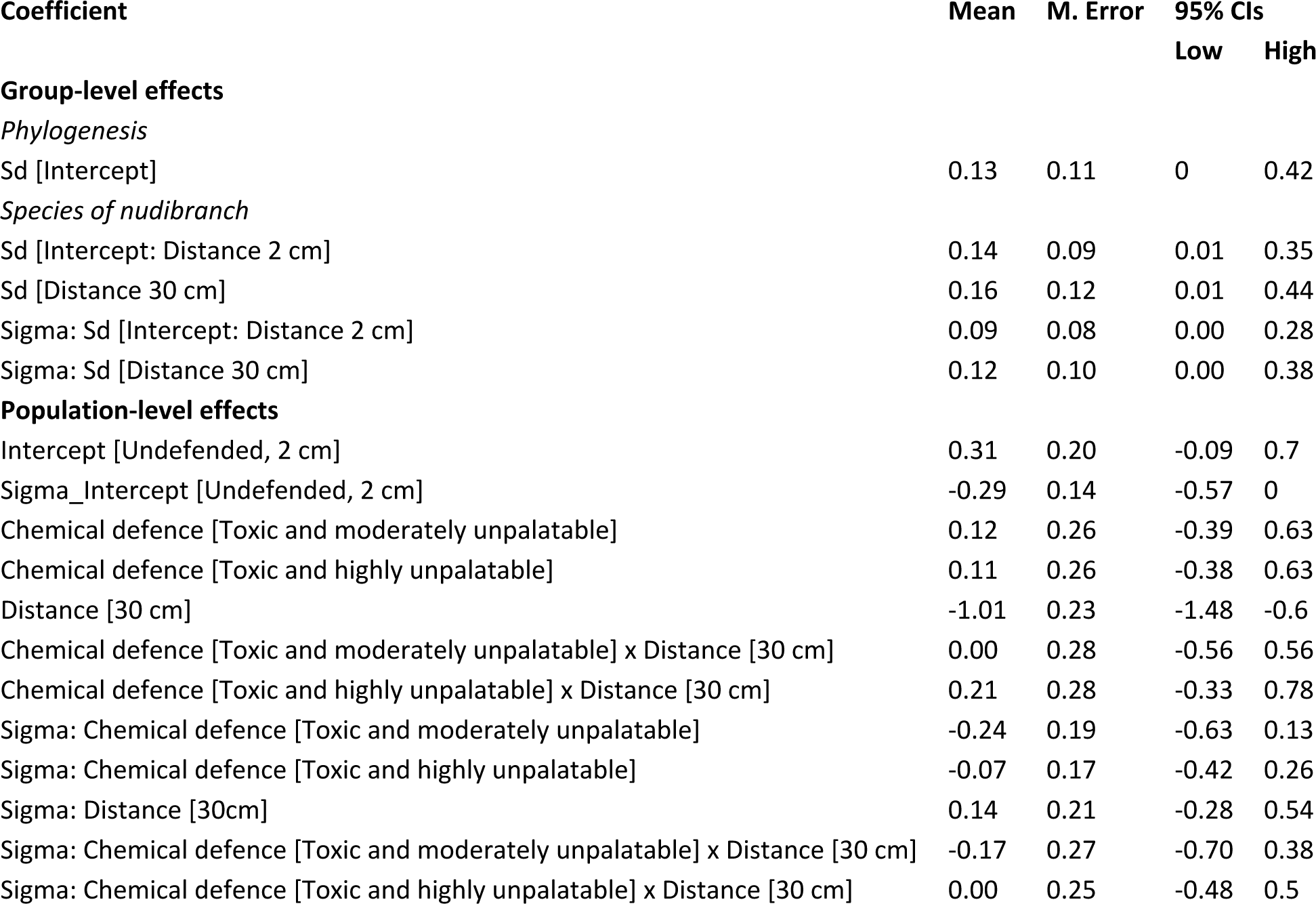
Coefficient estimates of the model investigating the scores for latent factor 2 between species of nudibranchs with different levels of chemical defences (N = 12, R^2 = 0.28). Estimates are based on a Student distribution with an identity link for the mean of the response distribution and a log link for its residual standard deviation (Sigma). The estimate is more likely to be non-zero when the credible intervals do not overlap with zero.

| Coefficient | Mean | M. Error | 95% CIs |  |
| --- | --- | --- | --- | --- |
|  |  |  | Low | High |
| Group-level effects |  |  |  |  |
| Phylogenesis |  |  |  |  |
| Sd [Intercept] | 0.13 | 0.11 | 0 | 0.42 |
| Species of nudibranch |  |  |  |  |
| Sd [Intercept: Distance 2 cm] | 0.14 | 0.09 | 0.01 | 0.35 |
| Sd [Distance 30 cm] | 0.16 | 0.12 | 0.01 | 0.44 |
| Sigma: Sd [Intercept: Distance 2 cm] | 0.09 | 0.08 | 0.00 | 0.28 |
| Sigma: Sd [Distance 30 cm] | 0.12 | 0.10 | 0.00 | 0.38 |
| Population-level effects |  |  |  |  |
| Intercept [Undefended, 2 cm] | 0.31 | 0.20 | -0.09 | 0.7 |
| Sigma_Intercept [Undefended, 2 cm] | -0.29 | 0.14 | -0.57 | 0 |
| Chemical defence [Toxic and moderately unpalatable] | 0.12 | 0.26 | -0.39 | 0.63 |
| Chemical defence [Toxic and highly unpalatable] | 0.11 | 0.26 | -0.38 | 0.63 |
| Distance [30 cm] | -1.01 | 0.23 | -1.48 | -0.6 |
| Chemical defence [Toxic and moderately unpalatable] x Distance [30 cm] | 0.00 | 0.28 | -0.56 | 0.56 |
| Chemical defence [Toxic and highly unpalatable] x Distance [30 cm] | 0.21 | 0.28 | -0.33 | 0.78 |
| Sigma: Chemical defence [Toxic and moderately unpalatable] | -0.24 | 0.19 | -0.63 | 0.13 |
| Sigma: Chemical defence [Toxic and highly unpalatable] | -0.07 | 0.17 | -0.42 | 0.26 |
| Sigma: Distance [30cm] | 0.14 | 0.21 | -0.28 | 0.54 |
| Sigma: Chemical defence [Toxic and moderately unpalatable] x Distance [30 cm] | -0.17 | 0.27 | -0.70 | 0.38 |
| Sigma: Chemical defence [Toxic and highly unpalatable] x Distance [30 cm] | 0.00 | 0.25 | -0.48 | 0.5 |

**Table S13.** Coefficient estimates of the model investigating the scores for latent factor 3 between species of nudibranchs with different levels of chemical defences (N = 12, R^2 = 0.07). Estimates are based on a Student distribution with an identity link for the mean of the response distribution and a log link for its residual standard deviation (Sigma). The estimate is more likely to be non-zero when the credible intervals do not overlap with zero.

| Coefficient | Mean | M. Error | 95% CIs |  |
| --- | --- | --- | --- | --- |
|  |  |  | Low | High |
| Group-level effects |  |  |  |  |
| Phylogenesis |  |  |  |  |
| Sd [Intercept] | 0.11 | 0.11 | 0 | 0.39 |
| Species of nudibranch |  |  |  |  |
| Sd [Intercept: Distance 2 cm] | 0.13 | 0.08 | 0.01 | 0.32 |
| Sd [Distance 30 cm] | 0.12 | 0.10 | 0.00 | 0.36 |
| Sigma: Sd [Intercept: Distance 2 cm] | 0.13 | 0.09 | 0.01 | 0.35 |
| Sigma: Sd [Distance 30 cm] | 0.12 | 0.10 | 0.00 | 0.36 |
| Population-level effects |  |  |  |  |
| Intercept [Undefended, 2 cm] | 0.22 | 0.18 | -0.14 | 0.58 |
| Sigma_Intercept [Undefended, 2 cm] | -0.49 | 0.19 | -0.87 | -0.1 |
| Chemical defence [Toxic and moderately unpalatable] | -0.23 | 0.24 | -0.69 | 0.27 |
| Chemical defence [Toxic and highly unpalatable] | -0.18 | 0.23 | -0.63 | 0.3 |
| Distance [30 cm] | -0.49 | 0.22 | -0.91 | -0.07 |
| Chemical defence [Toxic and moderately unpalatable] x Distance [30 cm] | 0.23 | 0.27 | -0.30 | 0.75 |
| Chemical defence [Toxic and highly unpalatable] x Distance [30 cm] | 0.17 | 0.25 | -0.32 | 0.66 |
| Sigma: Chemical defence [Toxic and moderately unpalatable] | -0.27 | 0.24 | -0.74 | 0.19 |
| Sigma: Chemical defence [Toxic and highly unpalatable] | -0.14 | 0.22 | -0.60 | 0.29 |
| Sigma: Distance [30cm] | 0.26 | 0.25 | -0.22 | 0.75 |
| Sigma: Chemical defence [Toxic and moderately unpalatable] x Distance [30 cm] | -0.11 | 0.32 | -0.75 | 0.51 |
| Sigma: Chemical defence [Toxic and highly unpalatable] x Distance [30 cm] | -0.27 | 0.30 | -0.87 | 0.32 |

**Table S14.**
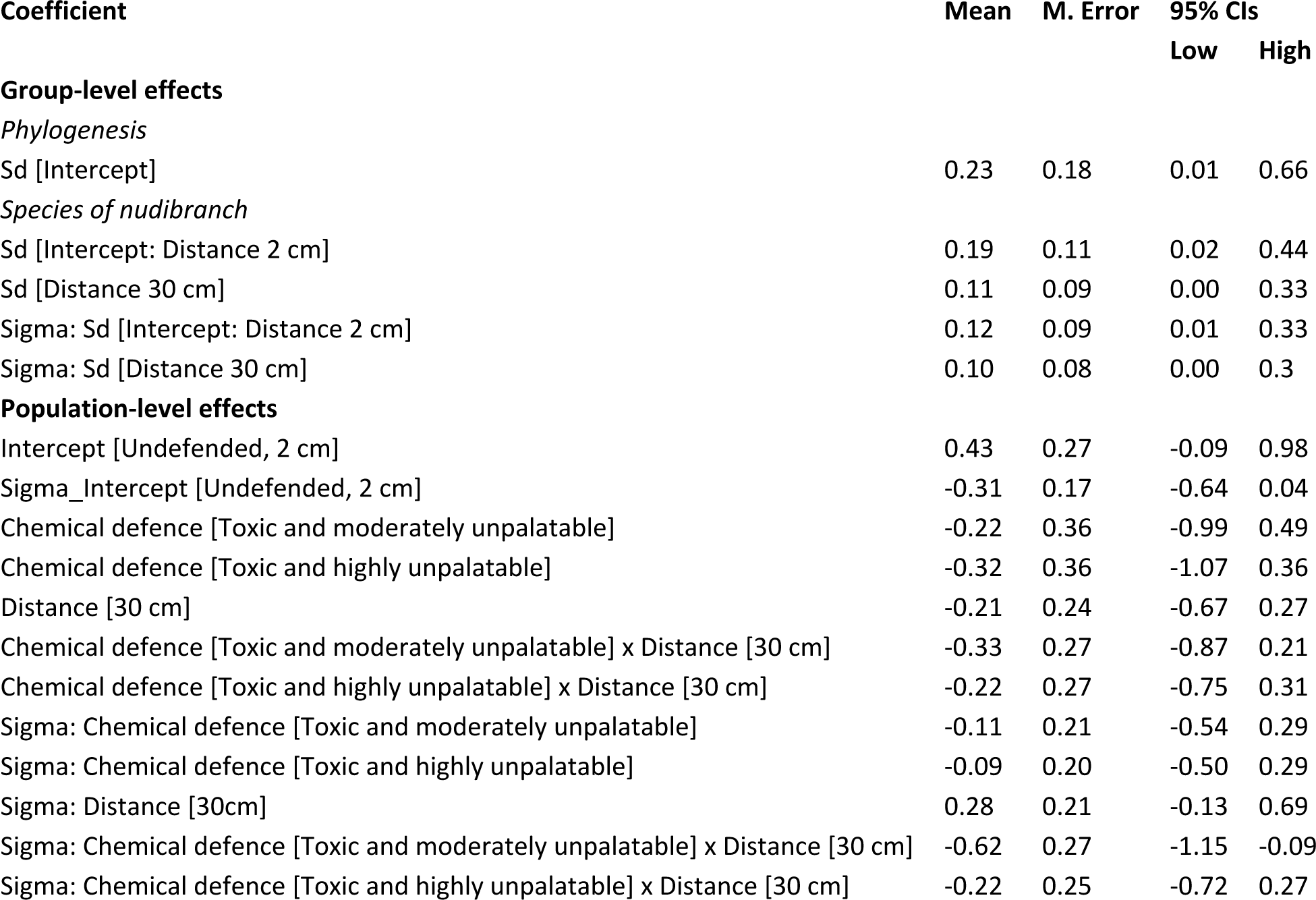
Coefficient estimates of the model investigating the scores for latent factor 4 between species of nudibranchs with different levels of chemical defences (N = 12, R^2 = 0.15). Estimates are based on a Student distribution with an identity link for the mean of the response distribution and a log link for its residual standard deviation (Sigma). The estimate is more likely to be non-zero when the credible intervals do not overlap with zero.

| Coefficient | Mean | M. Error | 95% CIs |  |
| --- | --- | --- | --- | --- |
|  |  |  | Low | High |
| Group-level effects |  |  |  |  |
| Phylogenesis |  |  |  |  |
| Sd [Intercept] | 0.23 | 0.18 | 0.01 | 0.66 |
| Species of nudibranch |  |  |  |  |
| Sd [Intercept: Distance 2 cm] | 0.19 | 0.11 | 0.02 | 0.44 |
| Sd [Distance 30 cm] | 0.11 | 0.09 | 0.00 | 0.33 |
| Sigma: Sd [Intercept: Distance 2 cm] | 0.12 | 0.09 | 0.01 | 0.33 |
| Sigma: Sd [Distance 30 cm] | 0.10 | 0.08 | 0.00 | 0.3 |
| Population-level effects |  |  |  |  |
| Intercept [Undefended, 2 cm] | 0.43 | 0.27 | -0.09 | 0.98 |
| Sigma_Intercept [Undefended, 2 cm] | -0.31 | 0.17 | -0.64 | 0.04 |
| Chemical defence [Toxic and moderately unpalatable] | -0.22 | 0.36 | -0.99 | 0.49 |
| Chemical defence [Toxic and highly unpalatable] | -0.32 | 0.36 | -1.07 | 0.36 |
| Distance [30 cm] | -0.21 | 0.24 | -0.67 | 0.27 |
| Chemical defence [Toxic and moderately unpalatable] x Distance [30 cm] | -0.33 | 0.27 | -0.87 | 0.21 |
| Chemical defence [Toxic and highly unpalatable] x Distance [30 cm] | -0.22 | 0.27 | -0.75 | 0.31 |
| Sigma: Chemical defence [Toxic and moderately unpalatable] | -0.11 | 0.21 | -0.54 | 0.29 |
| Sigma: Chemical defence [Toxic and highly unpalatable] | -0.09 | 0.20 | -0.50 | 0.29 |
| Sigma: Distance [30cm] | 0.28 | 0.21 | -0.13 | 0.69 |
| Sigma: Chemical defence [Toxic and moderately unpalatable] x Distance [30 cm] | -0.62 | 0.27 | -1.15 | -0.09 |
| Sigma: Chemical defence [Toxic and highly unpalatable] x Distance [30 cm] | -0.22 | 0.25 | -0.72 | 0.27 |

